# Non-canonical induction of autophagy increases adeno-associated virus type 2 (AAV2) transduction efficiency

**DOI:** 10.1101/2024.01.08.574727

**Authors:** Sereina O. Sutter, Sarah Jetzer, Anouk Lkharrazi, Sofia Pedersen, Elisabeth M. Schraner, Bernd Vogt, Hildegard Büning, Cornel Fraefel

## Abstract

Adeno-associated virus (AAV) serotypes infect a wide range of cell types, making this member of the parvovirus family a versatile tool in gene therapy. Infection as well as transduction is set in motion by means of specific receptors in conjunction with trafficking pathways, particularly endocytosis, a main cell entry pathway of non-enveloped viruses. Here, we report that efficacy of transduction is enhanced upon treating cells with hyperosmotic sucrose, a known blocker of clathrin-mediated endocytosis, through the non-canonical induction of autophagy. This mechanism of autophagy induction, however, is different from the previously reported AAV2-mediated induction of autophagy, which relies on a canonical, phosphoinositide 3-kinase class III (PI3K-III) complex-dependent pathway and appears to be dependent on the virus intrinsic secreted phospholipase A2 (sPLA_2_) domain, particularly its catalytic center activity.

**IMPORTANCE:** Adeno-associated virus (AAV) vectors are among the most frequently applied virus-based delivery vehicles for gene therapy. Lack of pathogenicity for humans, availability of a huge number of AAV serotypes differing in their cellular tropism, and the mainly episomal persistence of AAV vector genomes are clear advantages of these biological nanoparticles. By exploring non-pharmacological inducers of autophagy, we provide evidence for a potent and easy to apply strategy to significantly improve the efficacy of recombinant AAV-based gene delivery in hepatic and structural cells. Besides, our data also demonstrate the importance of autophagy for AAV2 infection and vector-mediated transduction in non-hepatic cells.

## INTRODUCTION

Adeno-associated virus type 2 (AAV2) is a small, non-pathogenic, helper virus-dependent parvovirus with a single-stranded (ss) DNA genome of approximately 5 kb, and is one the most frequently used viral vectors for gene therapy (1). In the absence of a helper virus, wild-type AAV2 can integrate its genome site-preferentially into the adeno-associated virus pre-integration site (AAVS1) on human chromosome 19 or persist in an episomal form in the nucleus (2, 3). Co-infection with a helper virus, such as adenovirus type 5 (AdV5), leads to entry into a lytic replication cycle, including the production of progeny virus particles (4). The AAV2 genome consists of two genes with multiple open reading frames (ORFs) flanked by 145 nt long inverted terminal repeats (ITRs) located on either side. The *rep* gene encodes the four non-structural Rep proteins, two of which are transcribed from the p5 and the p19 promoter, respectively. An alternative splice site regulates expression of the alternative transcripts, whereby the unspliced RNAs encode Rep78 and Rep52, whereas Rep68 and Rep40 are encoded by their corresponding spliced variant (5, 6). The three structural proteins VP1, VP2 and VP3, constituting the icosahedral capsid, are encoded by the *cap* gene. Furthermore, the *cap* gene encodes the assembly-activating protein (AAP) and the membrane-associated accessory protein (MAAP) by means of nested alternative ORFs (7, 8).

AAV serotypes have in common that they exhibit a broad cellular tropism (9). Referring to AAV2, the cellular receptors and host factors facilitating cell attachment, entry and intracellular trafficking include heparan sulfate proteoglycan, human fibroblast growth factor receptor 1, ⍺_V_β_5_ integrin, ⍺_5_β_1_ integrin (reviewed in (10)), the host factor KIAA0319L (synonymous AAVR) (11), and GPR108, a member of the G-protein coupled receptor family (12). Different entry pathways have been proposed for AAV2, including clathrin- and dynamin-dependent endocytosis or internalization supported by the Ras-related C3 botulinum toxin substrate 1 (Rac1), a small GTPase and a major effector of macropinocytosis (reviewed in (10)). However, internalization through clathrin-independent carriers (CLICs) and GPI-enriched endocytic compartments (GEECs) was also reported as a major endocytic infection route (13). It was shown that acidification in endocytic compartments and the activity of proteases trigger conformational changes of the AAV2 capsid, leading to the exposure of the N-terminal domain of the VP1 protein, known as VP1 unique region (VP1_u_). VP1_u_ contains a secreted phospholipase A2 homology domain (sPLA_2_) as well as nuclear localization signals that enable the endosomal escape of AAV2 and nuclear entry, respectively (14). Moreover, it was shown that AAV2 induces autophagy upon cell infection and that induction of the autophagic flux enhances AAV2 transduction in hepatocytes (15).

Autophagy is a highly conserved process in eukaryotes (16) required for the degradation of damaged organelles or proteins, invading pathogens or the maintenance of cellular homeostasis during various stress conditions (17). Three different types of autophagy exist, namely microautophagy, chaperone-mediated autophagy and macroautophagy (18). Microautophagy involves the engulfment of soluble constituents at the lysosomal membrane by autophagic tubes, which mediate invagination and vesicle scission into the lumen (19). This process leads to the delivery of cytoplasmic molecules into lysosomes, resulting in the degradation of the constituents. Chaperone-mediated autophagy depends on cytosolic chaperones, such as the cytosolic 70 kDa heat shock cognate protein (hsc70), which selectively delivers proteins to the lysosomal surface where the protein unfolds and crosses the lysosomal membrane (20). Macroautophagy, herein referred to as autophagy, is the most extensively studied type of this cellular process, where intracellular components are enclosed in a double-layered membrane vesicle called the autophagosome. Subsequently, the autophagosome fuses with lysosomes leading to the degradation of the enclosed components (21). The entire process is dependent on autophagy related genes (*Atg*) and proteins which sustain the core machinery of the autophagy pathway. The mammalian autophagic pathway can be considered as a flux, consisting of several steps: initiation, elongation, closure, maturation and degradation.

The Unc-51 like autophagy activating kinase-1 (ULK1) complex and the phosphoinositide-3-kinase class III (PI3K-III) complex are the key players in autophagy initiation. The ULK1 complex consists of the Unc-51 like kinase-1 (ULK1) and kinase-2 (ULK2) and is negatively regulated by the mammalian target of rapamycin (mTOR). During the initiation step of autophagy, mTOR, a serine/threonine protein kinase, or adenosine monophosphate activated protein kinase (AMPK) signals to the ULK1 complex, leading to inhibition or activation, respectively. AMPK has been shown to interact directly with ULK1 and to phosphorylate it in a nutrient-dependent manner (22). A classical scenario during starvation would be the inhibition of mTOR, which in turn leads to the activation of the ULK1 complex by directly phosphorylating ULK1 and Atg13, thereby leading to autophagy induction. The mTOR complex consists of 2 subunits, mTOR complex 1 (mTORc1) and mTOR complex 2 (mTORc2) (17), of which mTORc1 is primarily responsive to nutrient starvation.

Besides, the PI3K-III complex, consisting of the phosphatidylinositol 3-kinase (VPS34), beclin-1, the serine/threonine-protein kinase (VPS15) and Atg14L, has been proposed to be an additional regulator of autophagy induction and autophagosome formation (23). For the elongation step, the Atg12-Atg5-Atg16 complex and light chain 3 beta (LC3) conjugation systems are essential. As a first step, LC3 is processed by Atg4, leading to the production of LC3-I. Next, Atg7 and Atg3 support the conjugation of LC3-I with phosphatidylethanolamine (PE), resulting in the formation of LC3-II which further associates with the forming autophagosome. Closure refers to the last autophagosome formation step during which the phagophore, consisting of a single membrane, closes into a double membraned-organelle. This process is mediated through scission, during which the inner and outer autophagic membranes separate (24).

Maturation of autophagosomes requires its fusion with either a lysosomal or endosomal vesicle (25). Autophagosomes that fuse with endosomes are so-called amphisomes, while fusion of an autophagosome or amphisome with a lysosome results in an autolysosome. After lysosomal fusion with an autophagosome, LC3-II located on the outer membrane is cleaved off by Atg4 (18), whereas LC3-II located on the inner membrane is degraded by lysosomal enzymes (26). During the maturation process, autophagic vacuoles become acidic by fusion either with vesicles containing proton pumps such as endosomes, or with vesicles containing lysosomal membrane proteins and proton pumps, but no lysosomal enzymes (25).

In this study we addressed the following questions: firstly, whether AAV2 as a virus and as a vector can induce autophagy in non-hepatic cells, particularly fibroblasts; secondly, whether autophagy induction also enhances transduction efficiencies in these cells; and thirdly, whether AAV2-induced autophagy is canonical (ULK1 - and the PI3K-III complex-dependent), or non-canonical (independent of either the ULK1 complex or the PI3K-III complex).

Hence, by exploring (non-)pharmacological inducers of autophagy and the mechanism of AAV2-mediated induction of autophagy, we aim to investigate potent and easy to apply strategies to improve the efficacy of recombinant AAV vector-based gene delivery in hepatic and structural cells.

## RESULTS

### Autophagy induction enhances the transduction efficiency of recombinant AAV2 vectors

A recent study by Mardones et al. (27) reported that trehalose induces autophagy by inhibiting glucose transporters, resulting in the phosphorylation and activation of AMPK. As a result, AMPK activates the ULK1 complex independently of mTOR, leading to the induction of autophagy. Besides, sucrose, another non-reducing disaccharide, was shown to lead to the accumulation of lysosome-associated vacuoles (28) and to inhibit clathrin-mediated endocytosis (29) by preventing clathrin and adaptors from interacting (30). Moreover, it was demonstrated that the accumulation of sucrose containing vesicles, due to the absence of hydrolytic enzymes, resulted in the induction of autophagy (31).

To examine whether autophagy, induced by conventional drugs or simple sugars, impacts AAV2 transduction in hepatic, but also in non-hepatic cells, normal human fibroblasts (NHF) and human hepatocellular carcinoma (HepG2) cells were treated with sucrose or rapamycin (mTORc1 inhibitor) 1 h prior to infection with a recombinant AAV2 vector expressing a green fluorescent protein (GFP; rAAVGFP). As controls cells were either left untreated, or were treated with DMSO, chloroquine (lysosomal lumen alkalizer) or 3-methyladenine (3-Ma; PI3K-III inhibitor). As a vector dose, a multiplicity of infection of 2,000 genome containing particles (gcp) per cell (herein referred to as MOI) for NHF cells, and a MOI of 1,000 for HepG2 were chosen. At 24 hours post infection (hpi), transduction efficiency was assessed by fluorescence microscopy, showing an increased transduction efficiency in NHF (Fig. 1A) and HepG2 (Fig. 1B) cells treated with rapamycin or sucrose. Chloroquine and 3-Ma treatment, on the other hand, showed a significant decrease in transduction efficiency, indicating that autophagy induction by rapamycin, as well as sucrose treatment, increases AAV2 vector-mediated transduction of hepatic and non-hepatic cell types. A similar effect was observed upon the treatment of NHF cells with trehalose but not with the hydrolyzable sugars fructose and maltose (Fig. S1A). However, the observed sucrose-induced increase in transduction efficiency was not due to hyperosmolarity, as sorbitol did not have the same effect (Fig. S1B). Besides, sucrose treated cells infected with either a recombinant herpes simplex virus type 1 (HSV-C12) or adenovirus (Ad5GFP) showed a decrease in transduction efficiency or completely blocked transduction, respectively. As AdV is dependent on clathrin-mediated endocytosis for productive infection (32), the data indicates that sucrose is an efficient inhibitor of this pathway in NHF cells (Fig. S1C).

**Fig. 1:**
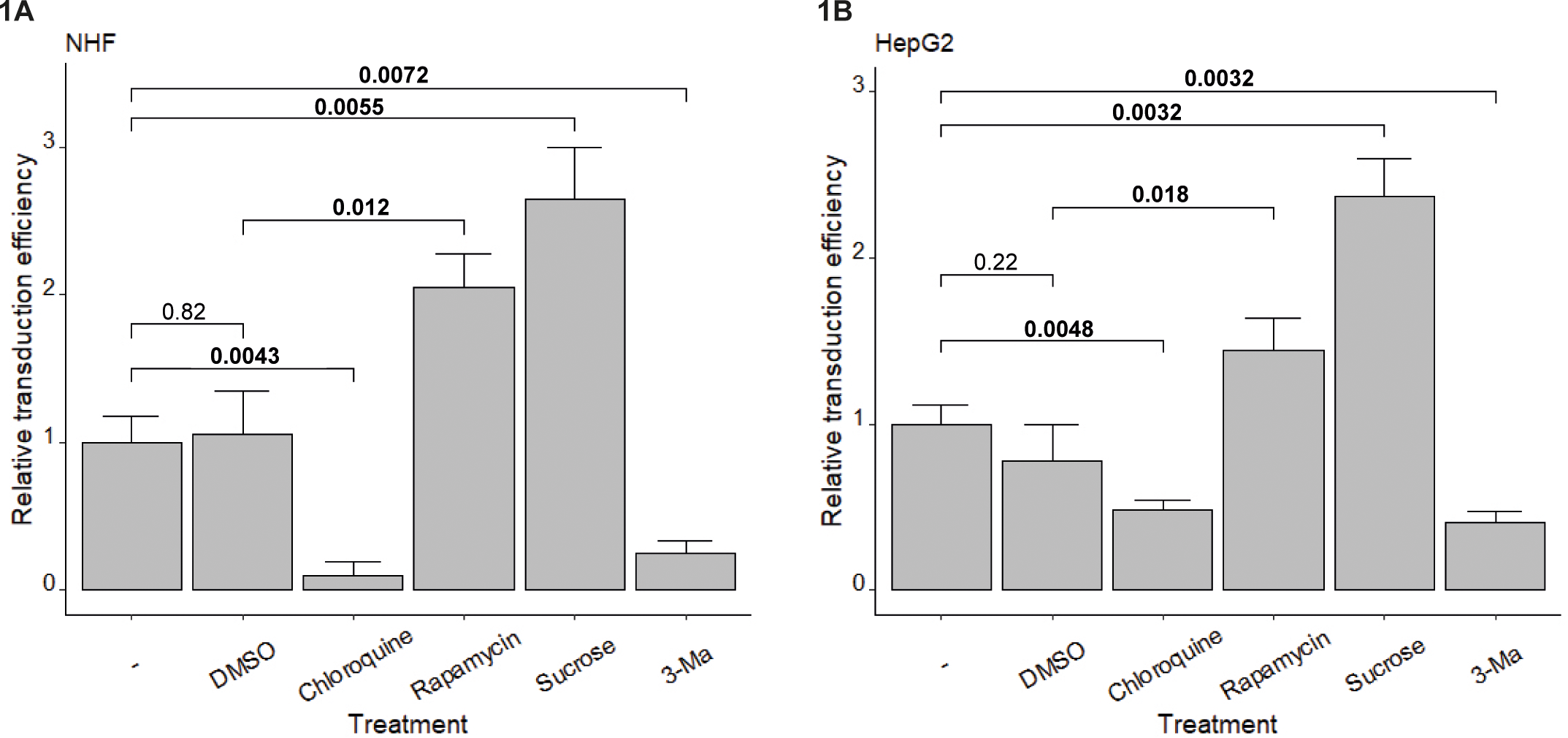
Autophagy induction enhances the transduction efficiency of recombinant AAV2 vectors. NHF (A) and HepG2 (B) cells were either untreated (-), or treated with DMSO, chloroquine, rapamycin, sucrose, or 3-Ma 1 h prior to infection with rAAVGFP (NHF: MOI 2,000, HepG2: MOI 1,000). At 24 hpi, transduction efficiency was determined by counting the numbers of GFP positive cells using an inverted fluorescence microscope. The graphs show mean values and standard deviations of the relative transduction efficiencies from triplicate experiments. p-values were calculated using Student’s t-test.

### Sucrose and AAV2 induce autophagy in non-hepatic cells

To confirm *de novo* induction of autophagic vesicle formation through AAV2 and sucrose in human primary fibroblasts, NHF cells and - as control - HepG2 cells were treated with rapamycin or sucrose, or infected with wild-type (wt) AAV2 (MOI 8,000) and compared to cells either left untreated, or treated with DMSO or chloriquine, respectively. In order to observe autophagic vesicle formation (autophagosomes, amphisomes, autolysosomes), cells were stained 5 hpi with the Cyto-ID autophagy detection reagent and monitored by confocal laser scanning microscopy (CLSM). The mean fluorescence intensity (MFI) of 70 cells per treatment was assessed with CellProfiler to determine the autophagy activity factor (AAF) in NHF (Fig. 2A) and HepG2 (Fig. 2B) cells. Cells treated either with chloroquine, rapamycin or sucrose, or infected with wtAAV2 showed an increased AAF compared to the negative controls. These results suggest that sucrose induces autophagic vesicle formation in NHF and HepG2 cells and in addition, it reveals that wtAAV2 induces autophagic vesicle formation not only in hepatic cells (15), but also in non-hepatic structural cells like fibroblasts.

**Fig. 2:**
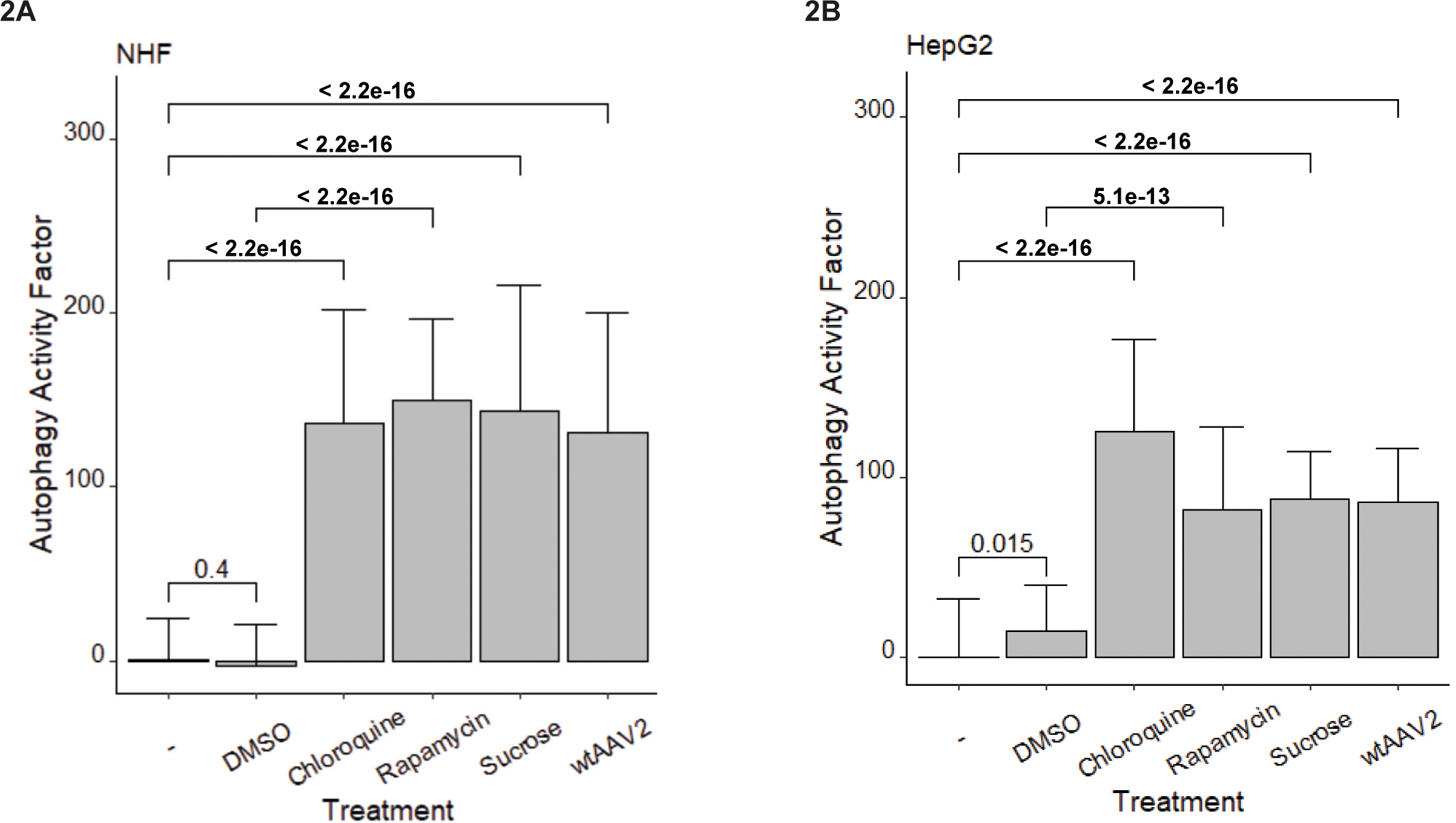
Sucrose and AAV2 induce autophagy in non-hepatic and hepatic cells. NHF (A) and HepG2 (B) cells were either untreated (-), or treated with DMSO, chloroquine, rapamycin, sucrose or infected with wtAAV2 (MOI 8,000). To observe autophagic vesicle formation, cells were stained with Cyto-ID 5 hpi, monitored by CLSM, and processed by image-based quantification using CellProfiler to determine the AAF. The graphs represent the mean values and standard deviations of the calculated AAF of 70 cells per treatment. p-values were calculated using Student’s t-test.

### Morphometric analysis of sucrose- and AAV2-induced autophagosome formation in human primary fibroblasts

As a further independent measure, autophagic vesicle formation was assessed by electron microscopy (EM). To this end, NHF cells were infected with wtAAV2 (MOI 20’000) in absence or presence of sucrose in the medium. At 5 hpi, the cells were fixed, sectioned and monitored by EM (Fig. 3A). Early and late autophagosomes in electron micrographs are indicated by white arrows. Nuclei (Nu), mitochondria (m) and Golgi (Gg) are indicated as well. The morphometric analysis of 25 cells per sample (Fig. 3B) revealed that sucrose treatment and wtAAV2 infection significantly increased autophagosome formation compared to untreated mock-infected cells. Moreover, sucrose treated and infected cells showed an even higher number of autophagosomes, indicating a cumulative effect of sucrose and wtAAV2 infection on autophagy induction in NHF cells.

**Fig. 3:**
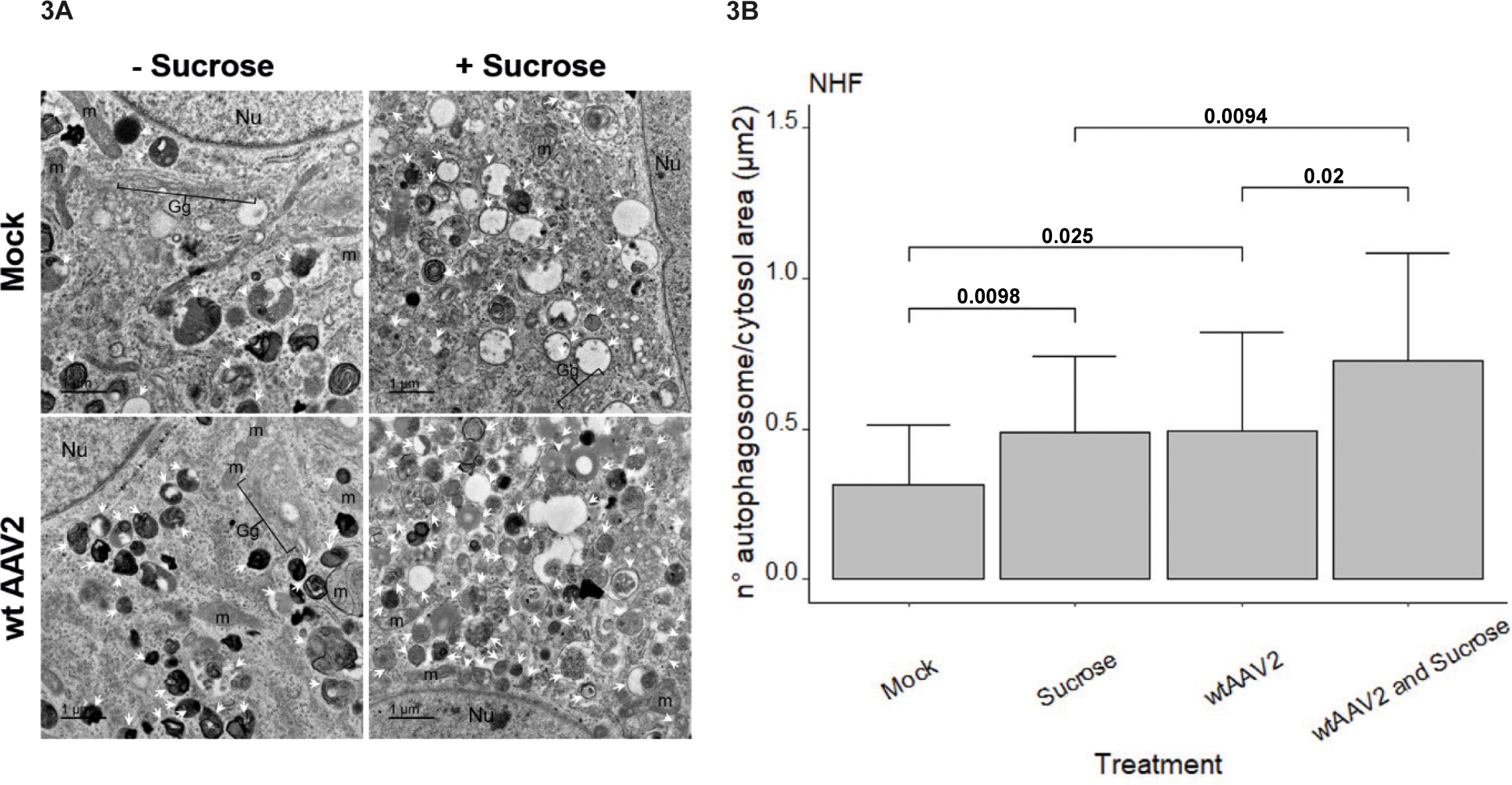
Sucrose and AAV2 increase the number autophagosomes in NHF cells. NHF cells were treated with sucrose 1 h prior to either mock-infection or infection with wtAAV2 (MOI 20,000). At 5 hpi, the cells were fixed, sectioned and processed for electron microscopy (A). Early and late autophagosomes are indicated by white arrows, nuclei (Nu), mitochondria (m) and Golgi (Gg) are indicated as well. The graph represents the mean values and standard deviations of number of autophagosomes per cytosol area in 25 cells per sample (B). p-values were calculated using Student’s t-test.

### Sucrose-induced autophagy is independent of the PI3K-III complex

3-Ma is widely used as an autophagy inhibitor as it blocks autophagosome formation at the initiation step by obstructing the kinase activity of VPS34 (17), an important subunit of the PI3K-III complex. Sucrose, on the other hand, was observed to increase autophagy, but it is not yet known whether sucrose induces autophagy through a canonical or a non-canonical pathway, *i.e.*, independently of the PI3K-III complex, as with trehalose (27).

To assess whether sucrose is able to induce autophagic vesicle formation independent of PI3K-III, *i.e.,* in conditions of PI3K-III inhibition, NHF (Fig. 4A) and HepG2 (Fig. 4B) cells were treated with either 3-Ma, sucrose, sucrose 1 h before 3-Ma, sucrose 1 h after 3-Ma or sucrose and 3-Ma combined. After 5 h, the cells were stained with Cyto-ID and monitored by CLSM. The MFI of 70 cells per treatment was determined by CellProfiler and the AAF was calculated. Both NHF and HepG2 cells showed an increased AAF upon sucrose treatment or treatment with different combinations of sucrose and 3-Ma. 3-Ma treatment alone, however, led to significantly reduced AAF, indicating that autophagic vesicle formation was inhibited. This data implies that sucrose-induced autophagy is not inhibited by 3-Ma treatment and strongly argues for a non-canonical pathway of autophagy induction.

**Fig. 4:**
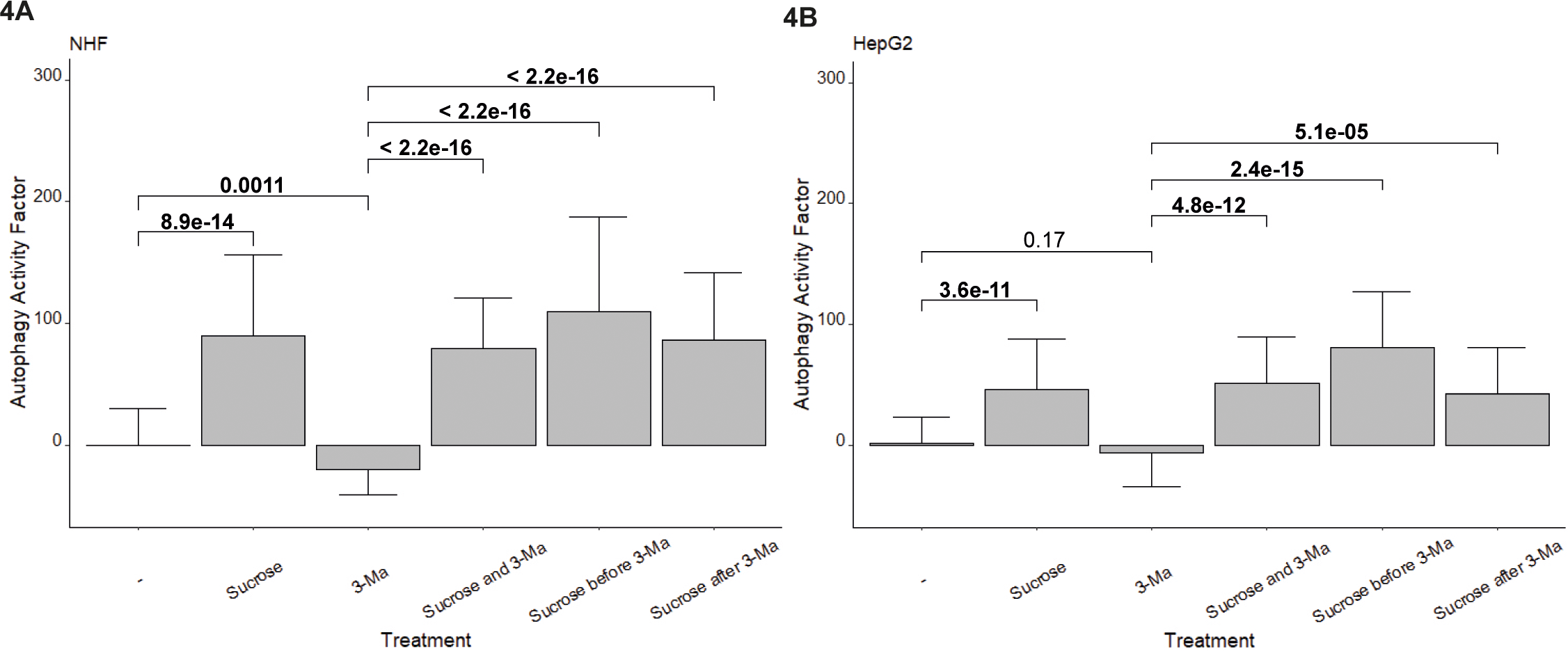
Sucrose-induced autophagy is independent of the PI3K-III complex. NHF (A) and HepG2 (B) cells were either untreated (-), or treated sucrose, 3-Ma, or sucrose and 3-Ma combined. To observe autophagic vesicle formation, cells were stained with Cyto-ID 5 hours post treatment, monitored by CLSM, and processed by image-based quantification using CellProfiler to determine the AAF. The graphs represent the mean values and standard deviations of the calculated AAF of 70 cells per treatment. p-values were calculated using Student’s t-test.

### Non-canonical induction of autophagy increases AAV2 vector transduction

As illustrated above, the combination of sucrose and 3-Ma treatment increased autophagy vesicle formation, likely by sucrose-mediated non-canonical autophagy induction. As canonical autophagy via the PI3K-III complex has been shown to enhance AAV2 vector-mediated cell transduction, the question arose whether an improved transduction is also observed in conditions of combined sucrose and 3-Ma treatment, and thus via the non-canonical induction pathway. To this end, NHF and HepG2 cells were treated with a combination of sucrose and 3-Ma 1 h prior to infection with rAAVGFP (NHF: MOI 2,000, HepG2: MOI 1,000). As control, cells were treated with sucrose, 3-Ma, or left untreated. At 24 hpi, transduction efficiency was assessed by fluorescence microscopy, showing an increased transduction efficiency in NHF (Fig. 5A) and HepG2 (Fig. 5B) cells upon treatment with either sucrose or sucrose and 3-Ma combined. 3-Ma treated cells showed a significant lower transduction rate compared to untreated cells, indicating that sucrose is able to overcome the inhibitory effect of 3-Ma by a non-canonical, PI3K-III complex-independent induction of autophagy, thereby leading to an increase of AAV2 vector transduction.

**Fig. 5:**
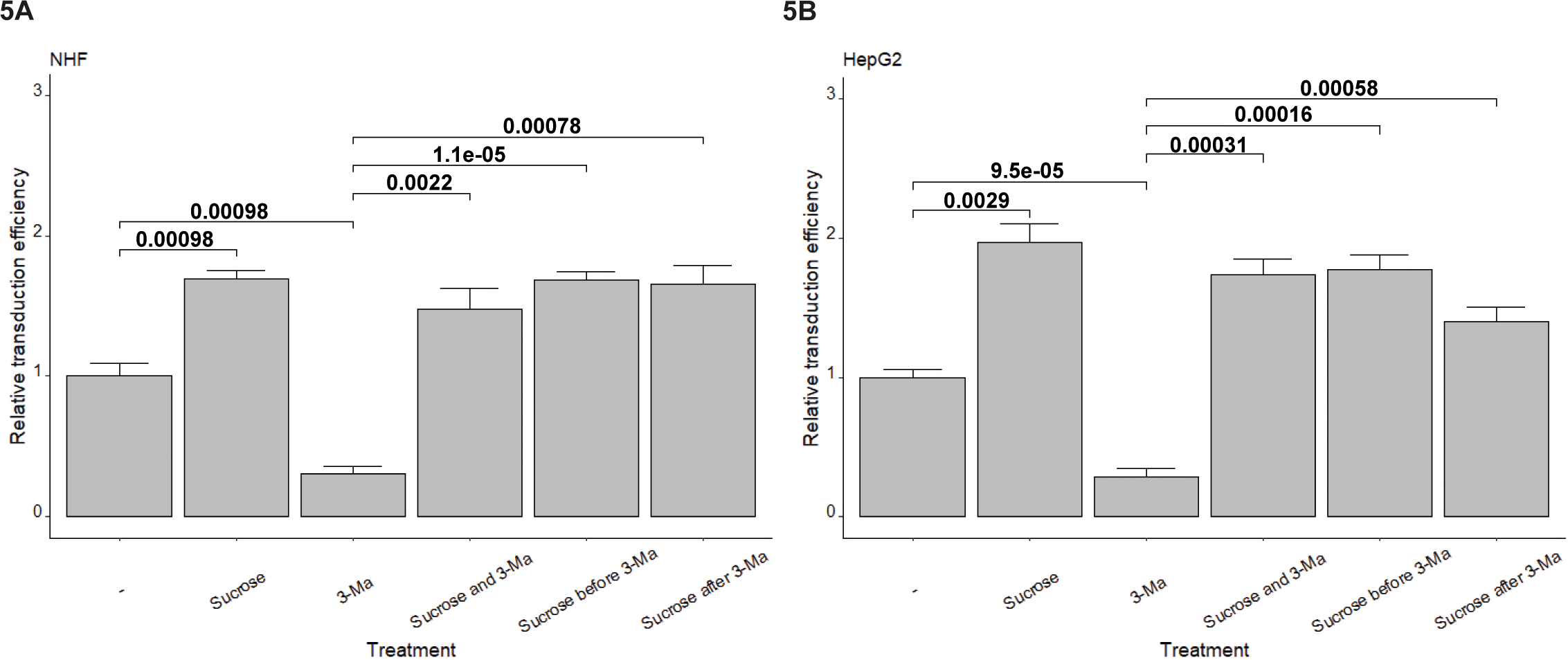
Non-canonical induction of autophagy increases AAV2 vector transduction. NHF (A) and HepG2 (B) cells were either untreated (-), or treated with sucrose, 3-Ma, or different combinations of sucrose and 3-Ma. After 1 h, cells were infected with rAAVGFP (NHF: MOI 2,000, HepG2: MOI 1,000). At 24 hpi, transduction efficiency was determined by counting the numbers of GFP positive cells using an inverted fluorescence microscope. The graphs show mean values and standard deviations of the relative transduction efficiencies from triplicate experiments. p-values were calculated using Student’s t-test.

### AAV2-induced autophagy induction depends on a functional PI3K-III complex

To analyze the effect of autophagy induction or inhibition on the capability of wtAAV2 to induce autophagic vesicle formation, NHF cells were either untreated, or treated with DMSO, rapamycin, sucrose, a combination of sucrose and 3-Ma, or 3-Ma alone 1 h prior to infection with wtAAV2 (MOI 8,000). At 5 hpi, cells were stained with Cyto-ID and monitored by CLSM. The AAF, determined by CellProfiler by assessing the MFI of 70 cells per treatment (Fig. 6), was increased in untreated wtAAV2 infected cells as well as in cells treated with either rapamycin, sucrose or the combination of sucrose and 3-Ma prior to infection. Treatment with 3-Ma alone 1 h prior to infection, however, strongly reduced the AAF compared to untreated wtAAV2 infected cells, indicating that wtAAV2 depends on a functional PI3K-III complex to induce autophagy in a canonical manner, confirming previous results obtained for hepatic cells (15).

**Fig. 6:**
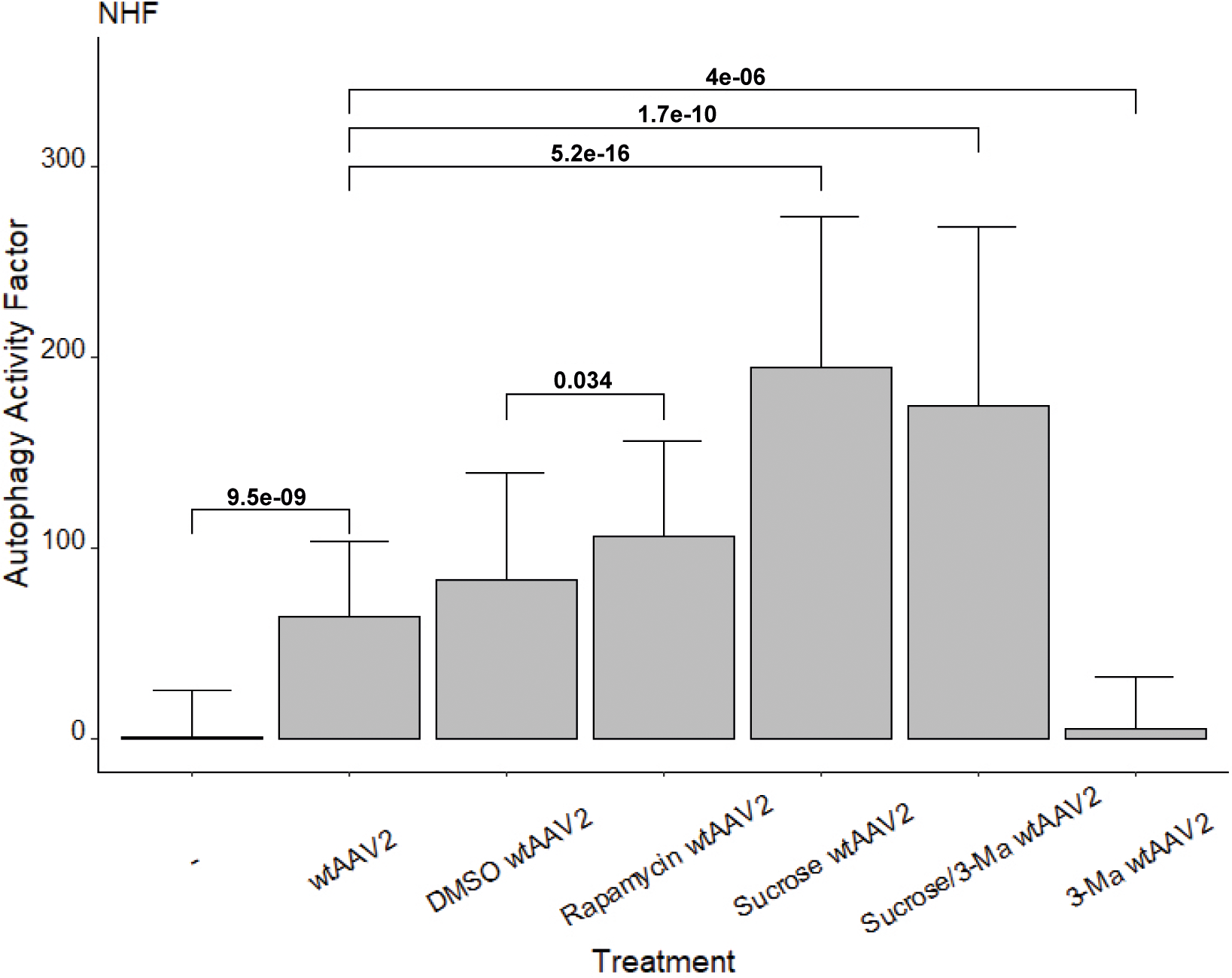
AAV2-induced autophagy induction depends on a functional PI3K-III complex. NHF cells were either untreated (-), or treated with DMSO, rapamycin, sucrose, sucrose and 3-Ma combined, or 3-Ma alone 1 h prior to infection with wtAAV2 (MOI 8,000). To observe autophagic vesicle formation, cells were stained with Cyto-ID 5 hpi, monitored by CLSM, and processed by image-based quantification using CellProfiler to determine the AAF. The graph represents the mean values and standard deviations of the calculated AAF of 70 cells per treatment. p-values were calculated using Student’s t-test.

### The post-transcriptional silencing of *VPS34* resulted in a minimal impact on AAV2 transduction

To further investigate the importance of the PI3K-III complex on AAV2 transduction, *VPS34* was post-transcriptionally silenced using small interfering RNA (siRNA). To this end, NHF cells were transfected with scrambled (scr) control siRNA or siRNA specifically targeting the coding sequence of *VPS34* (siVPS34 (33)). At 40 hours post transfection (hpt), the cells were treated with sucrose or trehalose 1 h prior to infection with rAAVGFP (MOI 2,000). As previously mentioned, trehalose has been shown to activate AMPK through the inhibition of glucose transporters, leading to ULK1 activation and autophagy induction independent of the PI3K-III complex (27). At 24 hpi, transduced cells were counted by fluorescence microscopy (Fig. 7A) and knock-down of *VPS34* (Fig. 7B) was confirmed on expression level by quantitative reverse transcription PCR (RT-qPCR). The post-transcriptional silencing of *VPS34*, either alone or in combination with sucrose or trehalose, resulted in a slight increase in AAV2 transduction compared to the scr control. This indicates that knock-down of *VPS34* might be beneficial for AAV2 vector-mediated transduction. Furthermore, scr control in combination with sucrose or trehalose did not show an increase in AAV2 transduction, which could be attributed to the lipofectamine-mediated induction of autophagy (34).

**Fig. 7:**
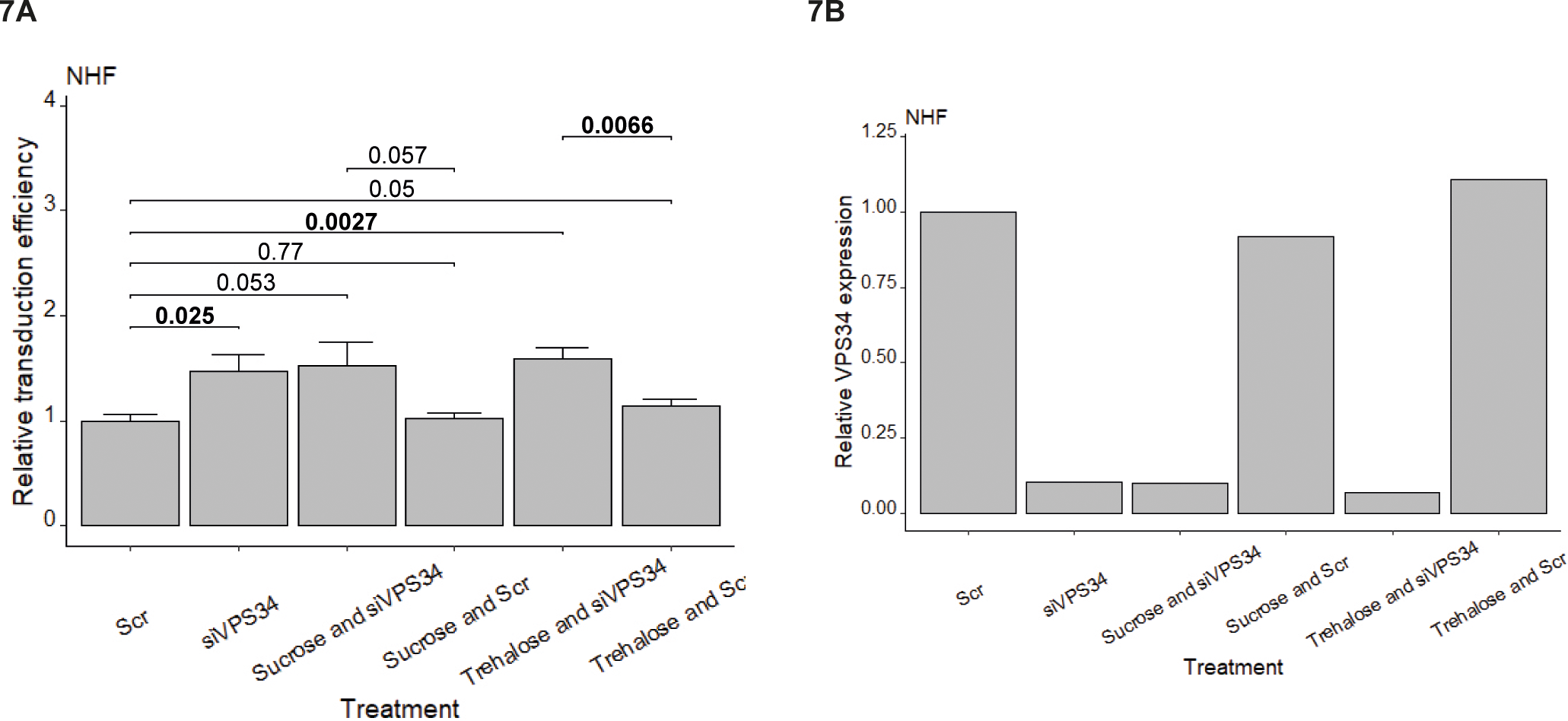
Knock-down of *VPS34* results in minimal impact on AAV2 vector transduction. NHF cells were transfected with scr control or *VPS34* targeting siRNA. At 40 hpt, cells were treated with sucrose or trehalose 1 h prior to infection with rAAVGFP (MOI 2,000). At 24 hpi, transduction efficiency was determined by counting the numbers of GFP positive cells using an inverted fluorescence microscope (A). The graphs show mean values and standard deviations of the relative transduction efficiencies from triplicate experiments. p-values were calculated using Student’s t-test. Knock-down of *VPS34* was confirmed on expression level by RT-qPCR (B).

### The ULK1 complex is essential for efficient AAV2 transduction

In addition to the PI3K-III complex, the ULK1 complex is another important element for the initiation of autophagy. As the PI3K-III complex was observed to be somewhat redundant for AAV2 transduction, the significance of the ULK1 complex in AAV2 transduction was assessed by post-transcriptional silencing of *Atg13*. To this end, NHF cells were either not transfected, or transfected with no, scr control, or siRNA specifically targeting the coding sequence *of Atg13*. At 40 hpt, the cells were treated with sucrose or trehalose 1 h prior to infection with rAAVGFP (MOI 2,000). At 24 hpi, transduced cells were assessed by fluorescence microscopy (Fig. 8A) and Western blot analysis (Fig. 8B). Post-transcriptional silencing of *Atg13*, either alone or in combination with sucrose or trehalose treatment, resulted in a significant decrease in AAV2 transduction. Treatment with sucrose and trehalose, alone or in combination with no or scr control siRNA, showed a significant increase in AAV2 transduction. These results indicate that Atg13 or a functional ULK1 complex, respectively, is essential for a successful transduction in NHF cells and that the sucrose-mediated increase in transduction efficiency depends on ULK1, likely through the ULK1-mediated induction of autophagy.

**Fig. 8:**
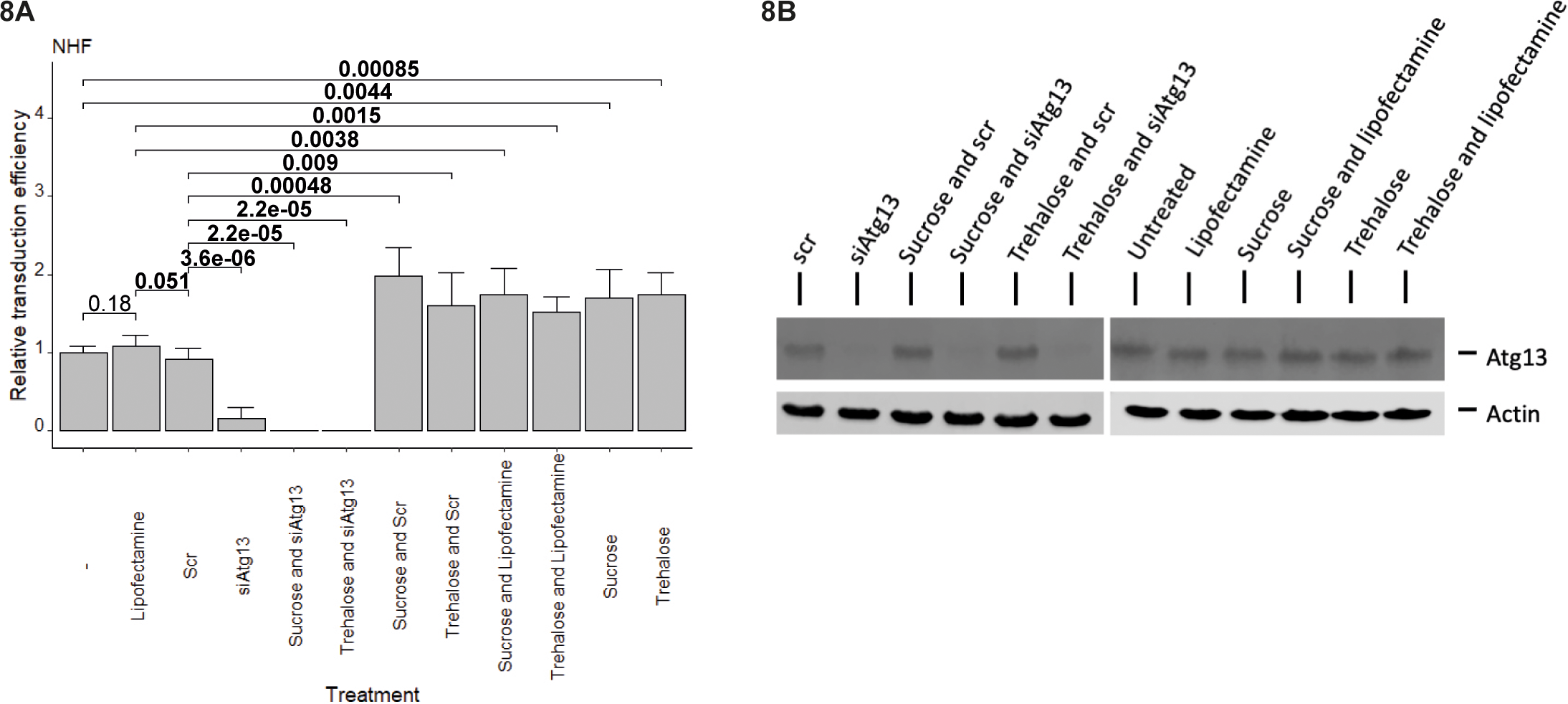
The ULK1 complex is essential for efficient AAV2 vector transduction. NHF cells were either not transfected (-), or transfected with no, scr control, or *Atg13* targeting siRNA. At 40 hpt, cells were treated with sucrose or trehalose 1 h prior to infection with rAAVGFP (MOI 2,000). At 24 hpi, transduction efficiency was determined by counting the numbers of GFP positive cells using an inverted fluorescence microscope (A). The graph shows mean values and standard deviations of the relative transduction efficiencies from triplicate experiments. p-values were calculated using Student’s t-test. Knock-down of *Atg13* was confirmed on protein level (B).

### The catalytic center of the PLA_2_ domain plays a pivotal function in endocytic vesicle escape, nuclear entry, and AAV2-mediated induction of autophagy

In addition to the nuclear localization sequences found in VP1 and VP2 (35, 36), a secreted phospholipase A_2_ (sPLA_2_) homology domain was identified (37), which is unique to VP1 and highly conserved among the *Parvoviridae*. This domain, however, is buried within the capsid interior and becomes externalized through pores located at the 5-fold symmetry axis during the passage of AAV as a virus or vector through the endosomal compartment (36, 38). The sequence similarity of both non-parvoviral and parvoviral sPLA_2_s is largely limited to the catalytic domain and to the calcium binding motif (39). Mutations in these regions drastically reduce enzymatic activity and viral infectivity, indicating a crucial role of the sPLA_2_ domain in the parvoviral life cycle (37, 40). Besides, it was reported that polyethyleneimine (PEI)-induced endosomal rupture or co-infection with AdV partially rescued the infectivity of a catalytic center sPLA_2_ mutant of the autonomously replicating parvovirus Minute Virus of Mice (MVM) and AAV2, demonstrating that the sPLA_2_ activity plays a role in rupturing the endosomal membrane (frequently referred to as lipolytic pore formation) to facilitate endosomal escape of incoming MVM and AAV2 particles (14, 41).

The detection of endocytic vesicle rupture, however, depends on galectins, which recognize the abnormal exposure of glycans to the cytosol after membrane damage, resulting either in a membrane damage response pathway or autophagy (reviewed in (42)). To address the question of whether the observed AAV2-mediated induction of autophagy results from sPLA_2_ activity-induced membrane rupturing, NHF cells were infected with AAV2 or a VP1 AAV2 mutant (^76^HD/AN) (40), which contains two mutated residues in the catalytic center of the PLA_2_ domain and is therefore deficient for endosomal escape. As control, cells were either left untreated, or treated with DMSO or rapamycin. At 5 hpi, the cells were stained with Cyto-ID and monitored by CLSM. The AAF, determined by CellProfiler by assessing the MFI of at least 50 cells per treatment (Fig. 9A), was increased in wtAAV2 infected cells as well as in cells treated with rapamycin. Infection with the VP1 AAV2 mutant (^76^HD/AN), however, strongly reduced the AAF compared to wtAAV2 infected cells, indicating that the catalytic center of the PLA_2_ domain plays a pivotal role in autophagy induction. Besides, AAV2- or VP1 AAV2 mutant infected cells were processed for immunofluorescence (IF) analysis to detect intact AAV2 capsids with fluorescence *in situ* hybridization (FISH) to visualize AAV2 genomes (Fig. 9B). The image-based quantification of AAV2 capsid counts of at least 50 cells or nuclei, respectively, at 3 (Fig. 9C) and 24 (Fig. 9D) hpi, showed a similar number in total cells, indicating that infection *per se* was not affected. The number of capsid counts in ^76^HD/AN-infected nuclei was strongly reduced compared to wild-type infected cells, indicating that the escape from endocytic vesicles and therefore nuclear entry was strongly impaired (this data and (14, 40)). Overall, these experiments imply that the catalytic center of the PLA_2_ domain plays not only a fundamental role in endocytic vesicle escape and nuclear entry, but also in the AAV2-mediated induction of autophagy.

**Fig. 9:**
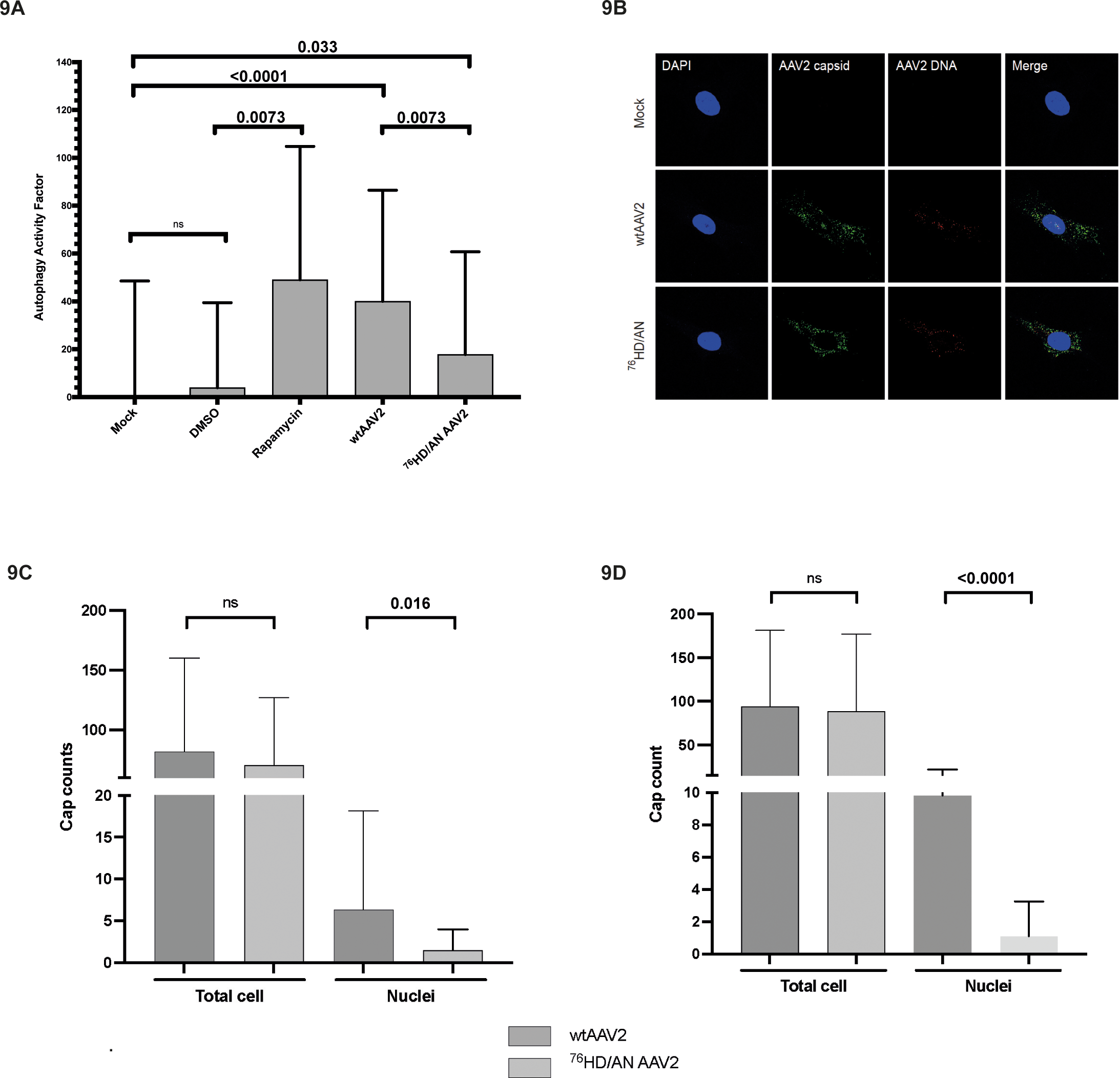
The VP1 AAV2 mutant (^76^HD/AN) induces autophagy less efficiently than wild-type AAV2. NHF cells were either untreated (Mock), or treated with DMSO, rapamycin, or infected with wtAAV2 or a VP1 AAV2 mutant (^76^HD/AN) at a MOI of 8,000. To observe autophagic vesicle formation, cells were stained with Cyto-ID 5 hpi, monitored by CLSM, and processed by image-based quantification using CellProfiler to determine the AAF (A). The graph represents the mean values and standard deviations of the calculated AAF of at least 50 cells per treatment. p-values were calculated using Student’s t-test. Wild-type AAV2 or a VP1 AAV2 mutant infected cells, were processed for IF-FISH and CLSM at 3 and 24 hpi (B). Intact capsids were stained using an antibody that detects a conformational capsid epitope (green). AAV2 DNA (red) was detected with an Alexa Fluor (AF) 647 labeled, amine-modified DNA probe that binds to the AAV2 genome. Nuclei were counterstained with DAPI (blue). Image-based quantification of AAV2 capsid counts in at least 50 cells or nuclei, respectively, was performed at 3 (9C) and 24 (9D) hpi using CellProfiler. p-values were calculated using an unpaired Student‘s t-test.

## DISCUSSION

Many viruses are taken up by receptor-mediated endocytosis and enter cells through the endocytic compartment. The fluent endosomal compartment provides a measure of protection from cytosolic innate immune sensors. However, viruses need to escape this compartment in order to reach their site of replication before they are either sorted to lysosomes or recycled back to the cell surface. Many enveloped viruses fuse their envelope with the endosomal membrane and release their capsid to the cytosol. Overall, this strategy is efficient and does not require any kind of membrane rupture. In contrast, non-enveloped viruses, with their hydrophilic capsids, have to cross membranes. In order to penetrate the endosomal membrane, most non-enveloped viruses undergo conformational changes and/or proteolytic processing that allow them to use membrane modulating factors. Nevertheless, not much is known about the mechanistic details of how non-enveloped viruses inflict membrane damage to cross endosomal membranes and even less is known about how they engage in the cellular response. In the case of parvoviruses, acidification in the endosome was shown to be a crucial step in order to induce conformational changes required for endosome penetration, as the drop in pH allows the deployment of the N-terminus of capsid protein VP1, possessing PLA_2_ activity (41, 43). This PLA_2_ activity seems to be essential for endosome penetration, most likely by transient and localized lipid modification (39, 41, 44). For example, canine parvovirus (CPV) entry allows the release of 3 kDa dextran, but not 10 kDa dextran, from endosomal compartments. The absence of co-release of larger molecules during parvovirus endosomal escape indicates that the virus-induced membrane rupture is limited and does not involve complete lysis (43). Specific to AAV2, it was demonstrated that the pH-dependent conformational changes of the AAV2 capsid results in the exposure of PLA_2_ and is important for AAV2 infection, partial uncoating (45), endosomal escape (14), and nuclear entry in particular (46). However, inflicting endocytic membrane damage results either in a galectin-mediated membrane damage response pathway or in the induction of autophagy. Autophagy is a fundamental process in eukaryotes (16), essential for survival under stresses such as starvation, invading pathogens as well as for cellular homeostasis (17). Intuitively, one would assume that the induction of autophagy is not favorable for a virus. However, in the case of AAV, not only does the viral infection *per se* induce autophagy, leading to increased transduction, but even more intriguingly, the application of pharmacological autophagy inducers results in significantly improved transgene expression *in vitro* and *in vivo* (15).

In this study, we report that not only pharmacological autophagy inducers, but also non-hydrolyzable sugars, such as sucrose, can induce autophagy in a PI3K-III complex independent way in both primary human fibroblasts and hepatocytes (summarized in Fig. 10).

**Fig. 10:**
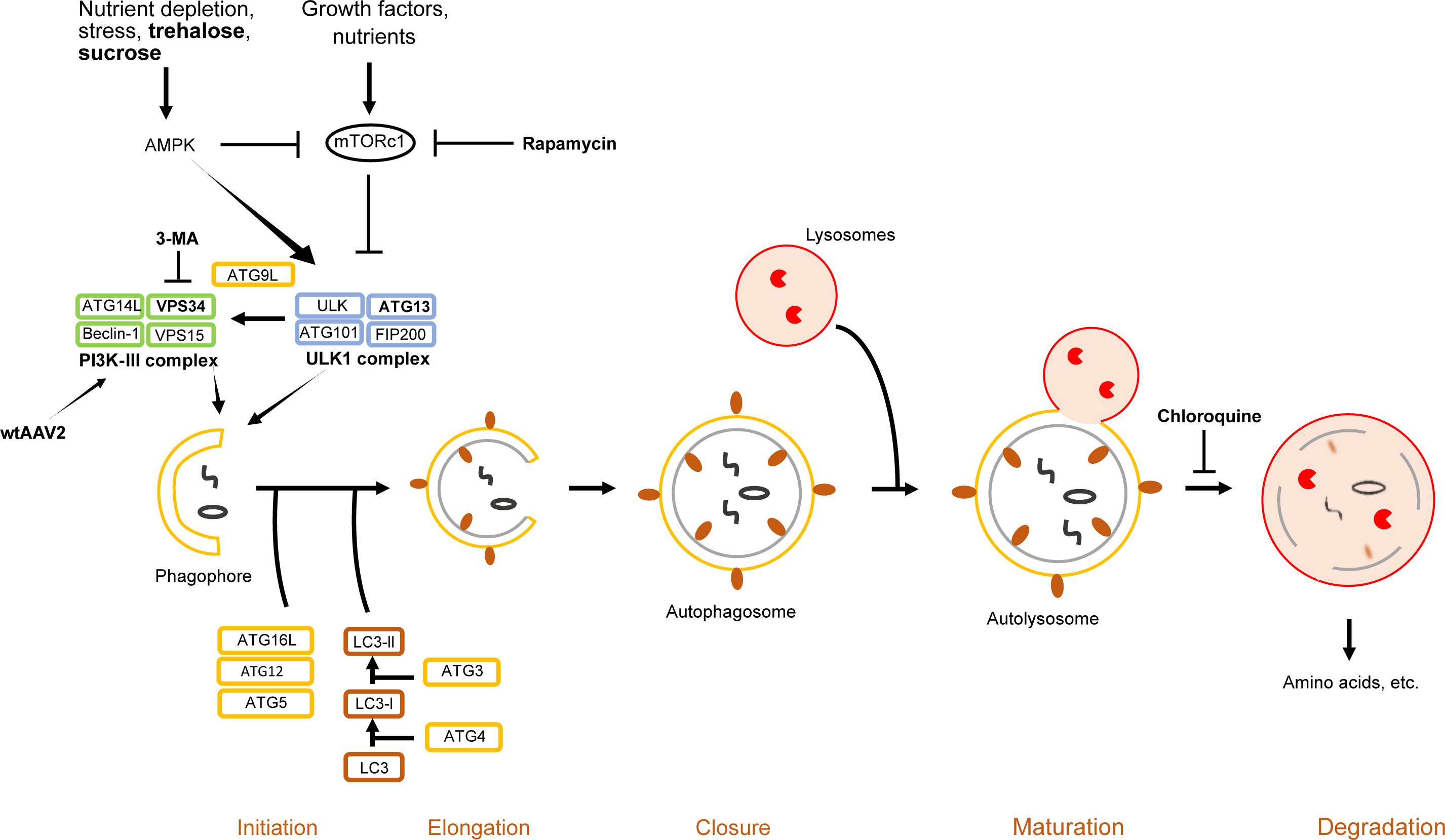
Schematic representation of the autophagic flux and its manipulation by (non-)pharmacological inducers, inhibitors, and AAV2 infection. Rapamycin induces autophagy by inhibiting mTORc1, thereby leading to the activation of the ULK1 complex. Non-hydrolyzable sugars, such as trehalose and sucrose, activate the ULK1 complex in an AMPK-dependent manner, resulting in the induction of autophagy. On the other hand, 3-Ma inhibits PI3K-III, resulting in the inhibition of autophagy. Besides, chloroquine inhibits the acidification of autolysosomes, which obstructs the autophagicflux. Wild-type AAV2 depends on a functional PI3K-III complex to induce autophagy in a canonical manner, in both, hepatic and non-hepatic cells. In addition, the ULK1 complex plays a pivotal function in efficient AAV2 vector transduction independent of the PI3K-III complex (not indicated in the figure). Essential components and drugs used in this study are indicated in bold.

Besides, we demonstrate that both pharmacological and non-pharmacological autophagy inducers strongly increase AAV2 vector-mediated transduction efficiency in human hepatocytes and fibroblasts, indicating that autophagy seems to play a pivotal function in AAV2 infection. It was shown that the pre-treatment of hepatocytes with torin 1 resulted in an increased level of intranuclear vector genomes (15), indicating that autophagic vesicles might be important for the retrograde transport of AAV2 virions towards the nucleus. Moreover, the vesicular acidification during autophagy, which occurs after fusion of autophagosomes with endosomes forming amphisomes and precedes the delivery of cytoplasmic cargo to lysosomes (25), might also favor different steps of the AAV life cycle (e.g., endosomal/amphisomal escape and nuclear entry). In contrast to the aforementioned findings, torin and rapamycin treatment failed to alter AAV2 transduction efficiency in HeLa cells (15). The same was observed in a study from Berry et al. (47), where neither the autophagy inducer nicardipine, nor the autophagy inhibitor spautin-1 altered AAV transduction in HeLa cells. These results indicate that not all cell lines are susceptible to the autophagy-mediated increase in AAV2 transduction efficiency. A possible explanation might be that in certain cancer cell lines, autophagy is already activated at a high level in order to escape ER stress-induced toxicity (48). The morphometric analysis of electron micrographs, however, showed that significantly more autophagosomes were formed upon sucrose treatment or AAV2 infection in human primary fibroblasts. The autophagosome formation increased even more when cells were first treated with sucrose prior to infection with wtAAV2, indicating a cumulative effect.

To investigate the pathway by which sucrose induces autophagy, human fibroblasts and hepatocytes were treated with a combination of sucrose and 3-Ma, revealing that sucrose is able to overcome the inhibitory effect of 3-Ma, indicating that the sucrose-mediated induction of autophagy is independent of the PI3K-III complex and therefore still able to increase AAV2 transduction efficiency in the presence of 3-Ma. Treatment of cells with 3-Ma prior to infection with wild-type AAV2, however, strongly reduced the induction of autophagy, indicating that the wild-type virus-mediated induction of autophagy depends on a functional PI3K-III complex in hepatic as well as non-hepatic cells. Nevertheless, the post-transcriptional silencing of *VPS34* showed a slight increase in AAV2 transduction efficiency compared to the scr control, indicating that the PI3K-III complex may not be essential for efficient AAV2 transduction. This observation is in contrast to the aforementioned image-based quantification of the AAF, which might be due to additional side effects of 3-Ma (49). Intriguingly, the post-transcriptional silencing of *Atg13* revealed that AAV2 transduction is significantly decreased even in the presence of sucrose or trehalose, indicating that the ULK1 complex is a crucial factor for efficient AAV2 transduction, at least in NHF cells. Moreover, our data indicate that sucrose, as well as trehalose (27), rely on the ULK1 complex in order to increase AAV2 transduction efficiency through autophagy induction. Furthermore, our image-based analysis, revealed that the catalytic center of the sPLA_2_ domain plays not only a fundamental role in endocytic vesicle escape and nuclear entry, but also in the wild-type AAV2-mediated induction of autophagy. Overall, our data demonstrate the importance of autophagy for AAV2 infection also in non-hepatic cells. Besides, by exploring non-pharmacological inducers of autophagy, we provide evidence for a potent strategy to significantly improve the efficacy of rAAV-based gene therapies in hepatic or structural cells, like human fibroblasts, respectively. And even though structural cells, like fibroblasts, are not a deliberate target in gene therapy, target tissue resident fibroblasts will become infected by AAV vectors.

## MATERIALS AND METHODS

### Cells

Normal human fibroblast (NHF) cells were kindly provided by X.O. Breakefield (Massachusetts General Hospital, Charlestown, MA, USA). NHF cells and HepG2 (ATCC HB-8065, American Type Culture Collection, Rockville, Md, USA were maintained in growth medium containing Dulbecco’s modified Eagle medium (DMEM) supplemented with 10% fetal bovine serum (FBS), 100 U/ml penicillin G, 100 μg/ml streptomycin, and 0.25 μg/ml amphotericin B (1% AB) at 37°C in a 95% air-5% CO_2_ atmosphere.

### Viruses

Wild-type (wt) AAV2 was produced by H. Büning (Hannover Medical School, Hannover, Germany). Recombinant AAVeGFP vectors of AAV serotype 2 were produced by transient transfection of 293T cells with pDG (50) and pAAVeGFP (kindly provided by M. Linden, King’s College London School of Medicine, London, UK) and purified by an iodixanol density gradient. Titers of genome-containing particles were determined by qPCR (51). The VP1 AAV2 mutant (^76^HD/AN) was constructed according to Girod et al. (40) and produced by the Viral Vector Facility (VVF) of the Neuroscience Center Zurich (ZNZ). Briefly, the ^76^HD/AN mutant construct was generated by mutating two key residues ^76^HD to ^76^AN using K-^76^HD/AN (5‘GCGGCCCTCGAGGCCAACAAAGCCTACGACCGG 3‘), L-^76^HD/AN (5‘CCGGTCGTAGGCTTTGTTGGCCTCGAGGGCCGC 3‘), psub-201 (52) containing the full-length AAV2 genome as template, and the QuikChange Site-Directed Mutagenesis Kit (Agilent Technologies). HSV-1 C12 (HSV-C12) is a recombinant HSV-1 strain SC16 containing a human cytomegalovirus (HCMV) IE1 enhancer/promoter-driven enhanced green fluorescent protein expression cassette in the US5 (gJ) locus and was kindly provided by S. Efstathiou (University of Cambridge, Cambridge, United Kingdom). Recombinant adenovirus (Ad5GFP) stocks were kindly provided by U. Greber (Department of Molecular Lice Sciences, University of Zurich, Switzerland).

### Treatment with autophagy inducers, inhibitors and sugars

NHF and HepG2 cells were treated with autophagy inducers, inhibitors, or sugars 1 h prior to mock-infection or infection with wtAAV2 or rAAVGFP. To this end, the cells were washed once with PBS and then treated with inducers, inhibitors, or sugars diluted in DMEM supplemented with 2% FBS and 1% AB (DMEM 2% FBS, 1% AB) according to the concentrations depicted in Table 1 and Table 2. The cells were then placed back in a humidified 95% air-5% CO_2_ incubator at 37°C.

**TABLE 1.**
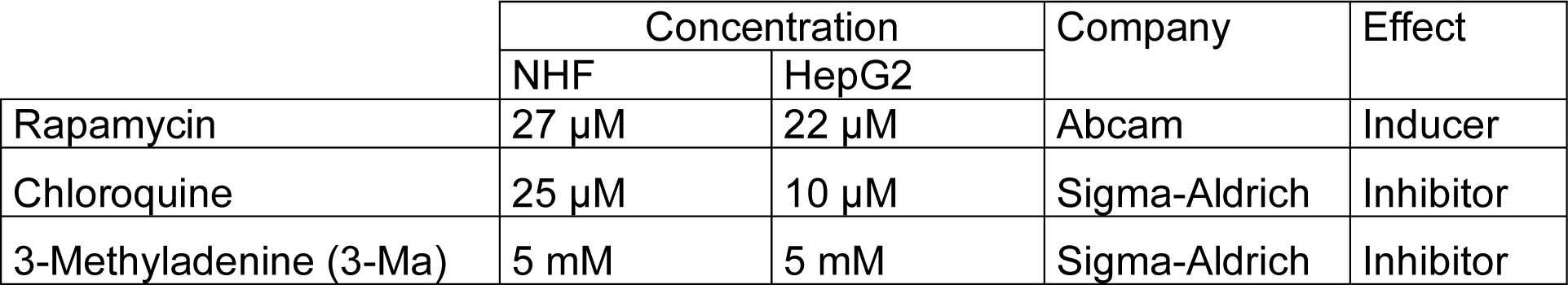
Drug concentrations used in NHF and HepG2 cells.

**TABLE 2.** Sugar and derivate concentrations used in NHF and HepG2 cells.

|  | Concentration |  | Company |
| --- | --- | --- | --- |
|  | NHF | HepG2 |  |
| Sucrose | 200 mM | 200 mM | Sigma-Aldrich |
| Trehalose | 200 mM | 200 mM | Sigma-Aldrich |
| Fructose | 200 mM | 200 mM | Sigma-Aldrich |
| Maltose | 200 mM | 200 mM | Sigma-Aldrich |
| Sorbitol | 200 mM | 200 mM | Sigma-Aldrich |

### Analysis of transduction efficiency

NHF or HepG2 cells were seeded into 24-well tissue culture plates at a density of 4×10^4^ or 5×10^4^ cells per well, respectively. The next day the cells were washed with PBS and either treated with DMSO, rapamycin, chloroquine, sucrose or 3-Ma according to Table 1 and Table 2. After 1 h, the cells were either mock-infected or infected with rAAVGFP at a MOI of 2,000 (NHF) or 1,000 (HepG2), or HSV-C12 (NHF: MOI 1), or Ad5GFP (NHF: MOI 5) in 250 μl of DMEM (0% FBS, 1% AB) pre-cooled to 4°C. The plates were first incubated for 30 min at 4°C to synchronize viral uptake and then incubated at 37°C in a humidified 95% air-5% CO_2_ incubator. At 24 hpi, the numbers of GFP-positive cells per well were determined with an inverted fluorescence microscope (Zeiss Axiovert S1000) and normalized to untreated cells.

### Cyto-ID assay

NHF and HepG2 cells were seeded on coverslips in a 24-well tissue culture plate at a density of 4×10^4^ or 5×10^4^ cells per well, respectively. Next, the cells were treated with autophagy inducers, inhibitors, or sucrose according to Table 1 and Table 2, or the cells were infected with either wtAAV2 (MOI 8,000) or the VP1 AAV2 mutant (^76^HD/AN: MOI 8,000). After 5 h, the cells were stained using the Cyto-ID autophagy detection kit according to the instructions of the manufacturer (Enzo Life Sciences). At least 50 cells per treatment were visualized using a confocal laser-scanning microscope (CLSM). To prevent cross talk between the channels for the different fluorochromes, all channels were recorded separately, and fluorochromes with longer wavelengths were recorded first. In order to measure the Cyto-ID mean fluorescence intensity (MFI) per cell, separate channel images of the Cyto-ID and the Hoechst staining were used as input images for a CellProfiler pipeline consisting of 5 modules. Each image was either assigned as “Cyto-ID” or “Hoechst” in order to define the module “IdentifyPrimaryObjects”. The following three modules (“ExpandOrShrinkObjects”, “MeasureObjectIntensity”, “ExportToSpreadsheet”) were used to determine the MFI per cell. The data obtained was further used to calculate the autophagy activity factor (AAF (53)) as follows:

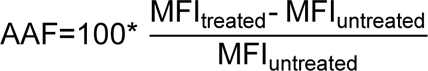

### RNA interference

9×10^4^ NHF cells per well were plated in 12-well tissue culture plates and transfected using lipofectamine RNAiMax transfection reagent (ThermoFisher scientific, Waltham, MA, USA) according to the manufacturer’s recommendations. The following RNAi oligonucleotides were used for the post-transcriptional silencing of *Atg13* (54) or *VSP34* (33), respectively: *Atg13* (CCA UGU GUG UGG AGA UUU CAC UUA A) and *VSP34* (CCC AUG AGA UGU ACU UGA ACG UAA UTT). Scr control siRNA was obtained from Santa Cruz Biotechnology (Control siRNA-A: sc-37007). At 40 hpt, the cells were washed once with PBS and treated with sucrose or trehalose according to Table 2 at 1 h prior to infection with rAAVGFP (MOI 2,000). Transduction efficiency was determined 24 hpi by counting numbers of GFP positive cells per well using an inverted fluorescence microscope (Zeiss Axiovert S1000) and normalized to untreated cells. Knock-down efficiency was assessed either by Western blot analysis or RT-qPCR.

### Antibodies

The following primary antibodies were used: anti-β-actin (Sigma-Aldrich A5316: dilution for Western blotting [WB], 1:5,000), anti-Atg13 (Cell signaling 13468: dilution for WB, 1:200), anti-AAV2 intact particle (A20, ProGen: dilution for Immunofluorescence [IF] 1:50). The following secondary antibodies were used: Alexa Fluor 488 goat anti-mouse IgG (Invitrogen A11001: dilution for IF 1:500), rabbit-anti mouse IgG-horseradish peroxidase (HRP: SouthernBiotech: dilution for WB 1:10,000) and goat-anti rabbit IgG-HRP (SouthernBiotech: dilution for WB 1:10,000).

### Western blot analysis

A total of 1.5×10^5^ NHF cells were seeded into 6-well tissue culture plates. The following day, the cells were washed with PBS and treated with sugars according to Table 2. After 24 h, the cells were trypsinized, washed once with PBS, and centrifuged for 5 min at 2000 x g and 4°C. The pellet was dissolved in 100 μl protein loading buffer (2.5% SDS, 5% β-mercaptoethanol, 10 % glycerol, 0.002% bromophenol blue, 62.5 mM Tris-HCl, pH 6.8), and then the samples were boiled for 10 min. Cell lysates were separated, depending on the molecular weight of the protein of interest, on 10% or 15% SDS-polyacrylamide gels and transferred to Protran nitrocellulose membranes (Whatman, Bottmingen, Switzerland). Membranes were blocked with PBS-T (PBS containing 0.3% Tween 20) supplemented with 5% non-fat dry milk for 1 h at room temperature (RT). Incubation with antibodies was carried out with PBS-T supplemented with 2.5% milk. Primary antibodies were incubated overnight at 4°C while secondary antibodies were incubated for 1 h at RT. Membranes were washed 3 times with PBS-T for 10 min after each antibody incubation step. HRP-conjugated secondary antibodies were detected by incubation with ECL (WesternBright^TM^ ECL-spray, Advansta Inc., Menlo Park, CA, USA) for 2 min. The membranes were exposed to chemiluminescence detection films (Roche Diagnistics, Rotkreuz, Switzerland). Detection of actin served as loading control for the lysate.

### RNA extraction

Cells were transfected, pre-treated and infected as described above. After 24 h, total RNA was extracted using the Direct-zol^TM^ RNA MiniPrep kit according to the instructions of the manufacturer (Zymo Research Corp, Irvine, CA, USA). DNA was digested by adding 8 µl of 10X DNase buffer, 5 µl DNase, 3 µl RNase-free water, and 64 µl RNA wash buffer and incubation for 15 min at 37°C. The samples were then purified according to the manufacturer’s (Zymo Research Corp) protocol. Quality and quantity of the extracted RNA was assessed using NanoDrop (Witec AG ND-1000 Spectrophotometer).

### Quantitative reverse transcription PCR (RT-qPCR)

To test the used primers, a standard RT-PCR (+/- RT reaction) with mock-infected NHF cells was performed. The cycling protocol started with a denaturation step of 3 min at 95°C, followed by 37 cycles of 30 sec at 94°C, 30 sec at 55°C and 1 min at 72°C followed by a final step of 10 min at 72°C. Subsequently, the reactions were analyzed on 1% agarose gels. Bands were expected between 100-200 bp, depending on the primer pair. The concentration and purity of RNA was determined by Qubit fluorometer analysis. To generate cDNA, the extracted RNA was reverse transcribed by using the reverse transcription system (Promega Corporation, Fitchburg, WI, USA). For this, the following components were mixed: 4 µl MgCl_2_, 2 µl 10X RT buffer, 2 µl dNTPs (10 mM), 0.5 µl of the RNase inhibitor RNAsin, 0.65 µl AMV RT, 1 µl random or Oligo(dT) primers, 1 µg RNA and RNase free H_2_O in a total volume of 20 µl. The mixture was incubated for 10 min at RT and 15 min at 42°C. For enzyme inactivation, the sample was incubated for 5 min at 95°C and then incubated on ice for 5 min. 2 µl of the cDNA were used for qPCR and the rest was stored at -20°C. For each reaction the following mixture was prepared: 0.25 μl forward primer (10 μM), 0.25 μl reverse primer (10 μM), 10 µl of SYBR Green PCR master mix and 7.5 µl ddH_2_O and transferred into a well of a Hard-Shell 96-well PCR plate (MicroAmp fast 96-well reaction plate). 2 µl of the appropriate cDNA was added and the 96-well plate was centrifuged for 1 min at 1000 x g and subsequently run at the standard 20 µl qPCR SYBR green program on QuantStudio 3 real-time system (Applied Biosystem, ThermoFisher Scientific, Waltham, MA, USA). The experiment was performed as technical triplicates for each primer pair for infected and non-infected samples and all conditions. *GAPDH* was used as housekeeping gene for the further normalization of the RT-qPCR raw data. Used primer sequences are listed in Table 3.

**TABLE 3.**
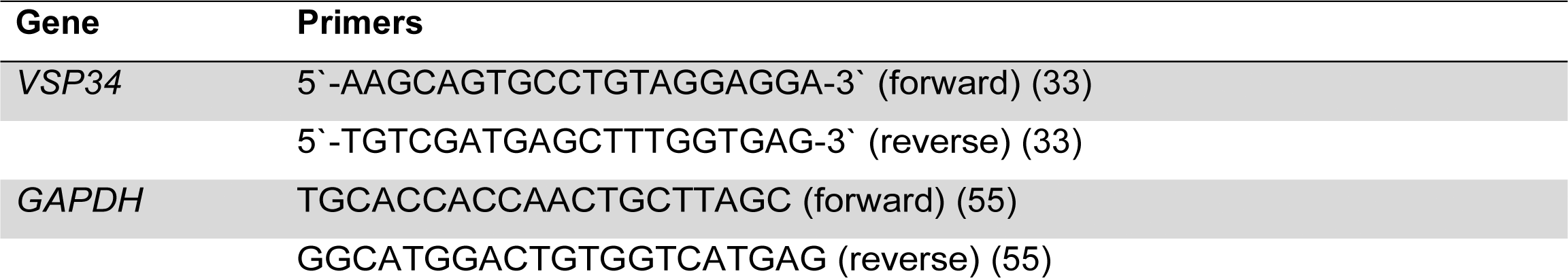
Primers used in this study.

### Combined multicolor immunofluorescence analysis and fluorescence *in situ* hybridization

FISH was performed essentially as described previously by Lux et al. (56). Briefly, a 3.9-kb DNA fragment containing the wtAAV2 genome without the inverted terminal repeats was amplified by PCR from plasmid pDG (50) using forward (5‘-CGGGGTTTTACGAGATTGTG-3‘) and reverse (5‘-GGCTCTGAATACACGCCATT-3‘) primers and the following conditions: 30 s at 95°C: 35 cycles of 10 s at 98°C, 15 s at 58°C, and 75 s at 72°C: and 10 min at 72°C. The PCR sample was then digested with DpnI to cut the residual template DNA and purified with the Pure Link PCR purification kit (Qiagen, Hilden, Germany). The DNA fragment was labeled with 5-(3-aminoallyl)dUTP by nick translation according to the manufacturer’s protocol (Ares DNA labeling kit, Molecular Probes, Eugene, OR, USA), and the incorporated dUTPs were labeled with amino-reactive Alexa Fluor 647 dye by using the same Ares DNA labeling kit.

4×10^4^ NHF cells were seeded onto coverslips (12-mm diameter: Glaswarenfabrik Karl Hecht GmbH & Co. KG, Sondheim, Germany) in 24-well tissue culture plates. The next day, the cells were treated with rapamycin, or infected with either wtAAV2 (MOI 8,000) or the VP1 AAV2 mutant (76HD/AN: MOI 8,000). At 3 or 24 hours after infection, the cells were washed with PBS, fixed for 30 min at RT with 2% PFA (in PBS), and washed again with PBS. The cells were then quenched for 10 min with 50 mM NH_4_Cl (in PBS), washed with PBS, permeabilized for 10 min with 0.2% Triton X-100 (in PBS), blocked for 10 min with 0.2% gelatin (in PBS) followed by two washing steps with PBS before blocking for 30 min in PBST (0.05% Tween 20 in PBS) supplemented with 3% BSA at 4°C. After antibody staining in PBST-BSA (3%, 25 μl/coverslip) for 1h at RT in the dark in a humidified chamber, the cells were washed three times for 5 min with PBST (0.1%), post-fixed with 2% PFA and blocked with 50 mM glycine in PBS for 5 min at RT. Hybridization solution (20 μl per coverslip) containing 1 ng/ml of the labeled DNA probe, 50% formamide, 7.3% (w/v) dextran sulfate, 15 ng/ml salmon sperm DNA, and 0.74x SSC (1x SSC is 0.15 M NaCl and 0.015 M sodium citrate) was denatured for 3 min at 95°C and shock-cooled on ice. The coverslips with the fixed and permeabilized cells facing down were placed onto a drop (20 μl) of the denatured hybridization solution and incubated overnight at 37°C in a humidified chamber (note that the cells were not denatured, as the AAV2 genome is present as ssDNA). The next day, the coverslips were washed three times with 2x SSC at 37°C, three times with 0.1x SSC at 60°C, and twice with PBS at RT. The cells were then embedded in ProLong Anti-Fade mountant (Molecular Probes, Eugene, OR, USA) containing 4‘,6-diamidino-2-phenylindole (DAPI) and imaged as midsections by confocal laser scanning microscopy (Leica SP8: Leica Microsystems, Wetzlar, Germany). To prevent cross talk between the channels for the different fluorochromes, all channels were recorded separately, and fluorochromes with longer wavelengths were recorded first. The resulting images were processed using Imaris V.7.7.2-V.9.6.0 (Bitplane, Oxford Instruments, Biplane AG, Zurich, Switzerland).

### Morphometric analysis autophagosome formation in human primary fibroblasts

1×10^6^ NHF cells were seeded into 10-cm tissue culture plates. The following day, the cells were treated with sucrose 1 h prior to mock-infection or infection with wtAAV2 (MOI 20,000). At 5 hpi, the cells were scraped off and transferred into a 15 ml conical centrifuge tube and pelleted for 5 min at 300 g. The cell pellet was resuspended in 2.5% glutaraldehyde in phosphate buffer, transferred to a 0.5 ml micro tube and centrifuged for 20 min at 3400 g. Afterwards the tip of the tube was carefully removed using a razor blade and the pellet was rinsed out in a glass vial using phosphate buffer and kept at 4°C. Next, the pellet was post-fixed in 1% OsO4 solution and dehydrated by an ascending ethanol series (70%, 80%, 96% for 10 min each, and 3 times in absolute ethanol for 10 min). Then, ethanol was exchanged by acetone in two steps, each lasting 15 minutes. Next, epon working solution was infiltrated at a 1:1 mix with acetone and incubated overnight. The next day, the pellet was transferred carefully into Easy Molds using a wooden stick and each mold was filled up with epon and placed into a desiccator for 6 h at RT. Polymerization was achieved by incubation at 60°C for 2.5 days followed by equilibration to RT. Next the samples were processed for ultrathin sectioning using an ultramicrotome. Next, the sections were collected on grids, stained with uranyl-acetate and lead citrate, and analyzed in a Philips CM 12 transmission electron microscope (Eindhoven, the Netherlands) equipped with a charge-coupled device (CCD) camera (Ultra-scan 1000, Gatan, Pleasanton, CA, USA) at an acceleration voltage of 100 kV.

### Image-based quantification and data analysis

For image-based quantification and data analysis, at least 50 individual cells per sample or condition were recorded and analyzed using different CellProfiler (V.2.2.0-V.4.0.7) pipelines. The output csv-files were further analyzed using Matlab (R2017a), GraphPad Prism 6 to 9, or R Studio 3.4.1. Depending on distribution frequency and standard deviation (SD), statistical analysis of individual experiments was either performed by unpaired Student‘s t-test or an unpaired t-test with Welch‘s correction (not assuming equal SDs). If not stated otherwise, each graph illustrates one representative experiment.

## ACKNOWLEDGMENTS

This work was supported by a grant (310030_184766 and 310030_212248) from the Swiss National Science Foundation to C.F.

## REFERENCES

1. Kuzmin DA, Shutova MV, Johnston NR, Smith OP, Fedorin VV, Kukushkin YS, Loo JCM van der, Johnstone EC. 2021. The clinical landscape for AAV gene therapies. Nat Rev Drug Discov 20:173–174.

2. Samulski RJ, Zhu X, Xiao X, Brook JD, Housman DE, Epstein N, Hunter LA. 1991. Targeted integration of adeno-associated virus (AAV) into human chromosome 19. The EMBO journal 10:3941–3950.

3. Sun X, Lu Y, Bish LT, Calcedo R, Wilson JM, Gao G. 2010. Molecular Analysis of Vector Genome Structures After Liver Transduction by Conventional and Self-Complementary Adeno-Associated Viral Serotype Vectors in Murine and Nonhuman Primate Models. Hum Gene Ther 21:750–761.

4. Buller RML, Janik JE, Sebring ED, Rose JA. 1981. Herpes Simplex Virus Types 1 and 2 Completely Help Adenovirus-Associated Virus Replication. J Virol 40:241– 247.

5. Srivastava A, Lusby EW, Berns KI. 1983. Nucleotide sequence and organization of the adeno-associated virus 2 genome. J Virol 45:555–564.

6. Laughlin CA, Westphal H, Carter BJ. 1979. Spliced adenovirus-associated virus RNA. Proc National Acad Sci 76:5567–5571.

7. Sonntag F, Kother K, Schmidt K, Weghofer M, Raupp C, Nieto K, Kuck A, Gerlach B, Bottcher B, Muller OJ, Lux K, Horer M, Kleinschmidt JA. 2011. The assembly-activating protein promotes capsid assembly of different adeno-associated virus serotypes. Journal of Virology 85:12686–12697.

8. Ogden PJ, Kelsic ED, Sinai S, Church GM. 2019. Comprehensive AAV capsid fitness landscape reveals a viral gene and enables machine-guided design. Science 366:1139–1143.

9. Asokan A, Schaffer DV, Samulski RJ. 2012. The AAV Vector Toolkit: Poised at the Clinical Crossroads. Mol Ther 20:699–708.

10. Nonnenmacher M, Weber T. 2012. Intracellular transport of recombinant adeno-associated virus vectors. Gene Ther 19:649–658.

11. Pillay S, Meyer NL, Puschnik AS, Davulcu O, Diep J, Ishikawa Y, Jae LT, Wosen JE, Nagamine CM, Chapman MS, Carette JE. 2016. An essential receptor for adeno-associated virus infection. Nature 530:108–112.

12. Dudek AM, Zabaleta N, Zinn E, Pillay S, Zengel J, Porter C, Franceschini JS, Estelien R, Carette JE, Zhou GL, Vandenberghe LH. 2020. GPR108 Is a Highly Conserved AAV Entry Factor. Mol Ther 28:367–381.

13. Nonnenmacher M, Weber T. 2011. Adeno-associated virus 2 infection requires endocytosis through the CLIC/GEEC pathway. Cell Host & Microbe 10:563–576.

14. Stahnke S, Lux K, Uhrig S, Kreppel F, Hösel M, Coutelle O, Ogris M, Hallek M, Büning H. 2011. Intrinsic phospholipase A2 activity of adeno-associated virus is involved in endosomal escape of incoming particles. Virology 409:77–83.

15. Hösel M, Huber A, Bohlen S, Lucifora J, Ronzitti G, Puzzo F, Boisgerault F, Hacker UT, Kwanten WJ, Klöting N, Blüher M, Gluschko A, Schramm M, Utermöhlen O, Bloch W, Mingozzi F, Krut O, Büning H. 2017. Autophagy Determines Efficiency of Liver-directed Gene Therapy with Adeno-associated Viral Vectors. Hepatology 10.1002/hep.29176.

16. King JS. 2012. Autophagy across the eukaryotes. Autophagy 8:1159–1162.

17. Yang Y, Hu L, Zheng H, Mao C, Hu W, Xiong K, Wang F, Liu C. 2013. Application and interpretation of current autophagy inhibitors and activators. Acta Pharmacol Sin 34:625–635.

18. Mizushima N, Yoshimori T, Levine B. 2010. Methods in Mammalian Autophagy Research. Cell 140:313–326.

19. Li W, Li J, Bao J. 2012. Microautophagy: lesser-known self-eating. Cell Mol Life Sci 69:1125–1136.

20. Kaushik S, Cuervo AM. 2012. Chaperone-mediated autophagy: a unique way to enter the lysosome world. Trends Cell Biol 22:407–417.

21. Feng Y, He D, Yao Z, Klionsky DJ. 2014. The machinery of macroautophagy. Cell Res 24:24–41.

22. Lee JW, Park S, Takahashi Y, Wang H-G. 2010. The Association of AMPK with ULK1 Regulates Autophagy. PLoS ONE 5:e15394.

23. Yuan H-X, Russell RC, Guan K-L. 2013. Regulation of PIK3C3/VPS34 complexes by MTOR in nutrient stress-induced autophagy. Autophagy 9:1983– 1995.

24. Knorr RL, Lipowsky R, Dimova R. 2015. Autophagosome closure requires membrane scission. Autophagy 11:2134–2137.

25. Eskelinen E-L. 2005. Maturation of Autophagic Vacuoles in Mammalian Cells. Autophagy 1:1–10.

26. Koukourakis MI, Kalamida D, Giatromanolaki A, Zois CE, Sivridis E, Pouliliou S, Mitrakas A, Gatter KC, Harris AL. 2015. Autophagosome Proteins LC3A, LC3B and LC3C Have Distinct Subcellular Distribution Kinetics and Expression in Cancer Cell Lines. PLoS ONE 10:e0137675.

27. Mardones P, Rubinsztein DC, Hetz C. 2016. Mystery solved: Trehalose kickstarts autophagy by blocking glucose transport. Sci Signal 9:fs2.

28. Cohn ZA, Ehrenreich BA. 1969. THE UPTAKE, STORAGE, AND INTRACELLULAR HYDROLYSIS OF CARBOHYDRATES BY MACROPHAGES. J Exp Med 129:201–225.

29. Heuser JE, Anderson RG. 1989. Hypertonic media inhibit receptor-mediated endocytosis by blocking clathrin-coated pit formation. J cell Biol 108:389–400.

30. Hansen SH, Sandvig K, Deurs B van. 1993. Clathrin and HA2 adaptors: effects of potassium depletion, hypertonic medium, and cytosol acidification. J cell Biol 121:61–72.

31. Higuchi T, Nishikawa J, Inoue H. 2015. Sucrose induces vesicle accumulation and autophagy. Journal of cellular biochemistry 116:609–617.

32. Nemerow GR, Pache L, Reddy V, Stewart PL. 2009. Insights into adenovirus host cell interactions from structural studies. Virology 384:380–388.

33. Itakura E, Kishi C, Inoue K, Mizushima N. 2008. Beclin 1 forms two distinct phosphatidylinositol 3-kinase complexes with mammalian Atg14 and UVRAG. Molecular biology of the cell 19:5360–5372.

34. Man N, Chen Y, Zheng F, Zhou W, Wen L-P. 2010. Induction of genuine autophagy by cationic lipids in mammalian cells. Autophagy 6:449–454.

35. Grieger JC, Snowdy S, Samulski RJ. 2006. Separate Basic Region Motifs within the Adeno-Associated Virus Capsid Proteins Are Essential for Infectivity and Assembly. J Virol 80:5199–5210.

36. Sonntag F, Bleker S, Leuchs B, Fischer R, Kleinschmidt JA. 2006. Adeno-associated virus type 2 capsids with externalized VP1/VP2 trafficking domains are generated prior to passage through the cytoplasm and are maintained until uncoating occurs in the nucleus. Journal of Virology 80:11040–11054.

37. Zádori Z, Szelei J, Lacoste M-C, Li Y, Gariépy S, Raymond P, Allaire M, Nabi IR, Tijssen P. 2001. A Viral Phospholipase A2 Is Required for Parvovirus Infectivity. Dev Cell 1:291–302.

38. Bleker S, Sonntag F, Kleinschmidt JA. 2005. Mutational Analysis of Narrow Pores at the Fivefold Symmetry Axes of Adeno-Associated Virus Type 2 Capsids Reveals a Dual Role in Genome Packaging and Activation of Phospholipase A2 Activity. J Virol 79:2528–2540.

39. Canaan S, Zádori Z, Ghomashchi F, Bollinger J, Sadilek M, Moreau ME, Tijssen P, Gelb MH. 2004. Interfacial Enzymology of Parvovirus Phospholipases A2 *. J Biol Chem 279:14502–14508.

40. Girod A, Wobus CE, Zádori Z, Ried M, Leike K, Tijssen P, Kleinschmidt JA, Hallek M. 2002. The VP1 capsid protein of adeno-associated virus type 2 is carrying a phospholipase A2 domain required for virus infectivity. The Journal of general virology 83:973–978.

41. Farr GA, Zhang L, Tattersall P. 2005. Parvoviral virions deploy a capsid-tethered lipolytic enzyme to breach the endosomal membrane during cell entry. Proc Natl Acad Sci 102:17148–17153.

42. Daussy CF, Wodrich H. 2020. “Repair Me if You Can”: Membrane Damage, Response, and Control from the Viral Perspective. Cells 9:2042.

43. Suikkanen S, Antila M, Jaatinen A, Vihinen-Ranta M, Vuento M. 2003. Release of canine parvovirus from endocytic vesicles. Virology 316:267–280.

44. Dorsch S, Liebisch G, Kaufmann B, Landenberg P von, Hoffmann JH, Drobnik W, Modrow S. 2002. The VP1 Unique Region of Parvovirus B19 and Its Constituent Phospholipase A2-Like Activity. J Virol 76:2014–2018.

45. Sutter SO, Lkharrazi A, Schraner EM, Michaelsen K, Meier AF, Marx J, Vogt B, Büning H, Fraefel C. 2022. Adeno-associated virus type 2 (AAV2) uncoating is a stepwise process and is linked to structural reorganization of the nucleolus. Plos Pathog 18:e1010187.

46. Bartlett JS, Wilcher R, Samulski RJ. 2000. Infectious entry pathway of adeno-associated virus and adeno-associated virus vectors. Journal of Virology 74:2777– 2785.

47. Berry GE, Asokan A. 2016. Chemical Modulation of Endocytic Sorting Augments Adeno-associated Viral Transduction. The Journal of biological chemistry 291:939– 947.

48. Ogata M, Hino S, Saito A, Morikawa K, Kondo S, Kanemoto S, Murakami T, Taniguchi M, Tanii I, Yoshinaga K, Shiosaka S, Hammarback JA, Urano F, Imaizumi K. 2006. Autophagy Is Activated for Cell Survival after Endoplasmic ReticulumStress. Mol Cell Biol 26:9220–9231.

49. Huotari J, Helenius A. 2011. Endosome maturation. EMBO J 30:3481–3500.

50. Grimm D, Kern A, Rittner K, Kleinschmidt JA. 1998. Novel Tools for Production and Purification of Recombinant Adenoassociated Virus Vectors. Hum Gene Ther 9:2745–2760.

51. Grieger JC, Choi VW, Samulski RJ. 2006. Production and characterization of adeno-associated viral vectors. Nat Protoc 1:1412–1428.

52. Samulski RJ, Chang LS, Shenk T. 1987. A recombinant plasmid from which an infectious adeno-associated virus genome can be excised in vitro and its use to study viral replication. J Virol 61:3096–3101.

53. Chan LL-Y, Shen D, Wilkinson AR, Patton W, Lai N, Chan E, Kuksin D, Lin B, Qiu J. 2012. A novel image-based cytometry method for autophagy detection in living cells. Autophagy 8:1371–1382.

54. Hosokawa N, Hara T, Kaizuka T, Kishi C, Takamura A, Miura Y, Iemura S, Natsume T, Takehana K, Yamada N, Guan J-L, Oshiro N, Mizushima N. 2009. Nutrient-dependent mTORC1 Association with the ULK1–Atg13–FIP200 Complex Required for Autophagy. Mol Biol Cell 20:1981–1991.

55. Vandesompele J, Preter KD, Pattyn F, Poppe B, Roy NV, Paepe AD, Speleman F. 2002. Accurate normalization of real-time quantitative RT-PCR data by geometric averaging of multiple internal control genes. Genome biology 3:RESEARCH0034.

56. Lux K, Goerlitz N, Schlemminger S, Perabo L, Goldnau D, Endell J, Leike K, Kofler DM, Finke S, Hallek M, Büning H. 2005. Green fluorescent protein-tagged adeno-associated virus particles allow the study of cytosolic and nuclear trafficking. Journal of Virology 79:11776–11787.

